# Monoaminergic variation, cortical structure, and disaster trauma interact to shape emotional vulnerability

**DOI:** 10.64898/2026.01.29.702665

**Authors:** Daiki X. Sato, Takashi Makino, Kentaro Katahira, Zhiqian Yu, Hiroaki Tomita, Shunji Mugikura, Kengo Kinoshita, Masakado Kawata

## Abstract

Emotional vulnerability arises from the interplay of genetic variation, cortical network capacity, and subjective processing of adverse experiences, yet these components are rarely examined together in population-scale datasets. Monoaminergic signaling offers an opportunity for understanding such interactions because variation in related genes shapes sensitivity to environmental inputs. Among them, the vesicular monoamine transporter 1 (*VMAT1*) Thr136Ile variant is notable for its functional effects and its long-term maintenance at intermediate frequencies, a pattern consistent with context-dependent selection. These properties make it an informative marker for investigating how genetic sensitivity interacts with neurobiological and experiential factors to shape affective functioning. Using data from up to 9,625 participants in the Tohoku Medical Megabank Project, a cohort established after the 2011 Great East Japan Earthquake, we investigated how monoaminergic variation, cortical morphology, and traumatic memory shape affective functioning. We found associations between the Thr136Ile variant and negative affectivity and depressive symptoms, consistent with prior reports of heightened emotional reactivity associated with the 136Thr allele. However, magnitude of these effects was substantially amplified in specific experiential and neurobiological contexts. Subjective trauma sensitivity, capturing the discrepancy between earthquake disruption and current traumatic memory, displayed a genotype-dependent association with depressive symptoms, and this relationship was strongest when parietal or insular cortical surface area was smaller. Together, these results identify a multilevel pathway through which monoaminergic variation contributes to emotional vulnerability by interacting with both cortical network capacity and trauma processing. The context-dependent influence of monoaminergic variation may further contribute to maintenance of affective diversity in human populations.

## Introduction

Emotion and stress regulation rely on large-scale neural systems that integrate sensory, cognitive, and interoceptive information. Monoaminergic neurotransmitters shape corticolimbic circuits involved in threat detection, valuation, and regulatory control (Robbins and Arnsten 2009; Cools and D’Esposito 2011; Grace 2016). These systems influence momentary emotional responses as well as long-term stress sensitivity, making them key contributors to individual differences in affective functioning (McGaugh 2004; Joëls and Baram 2009). Such variability reflects genetic influence, environmental experience, and neural structural differences (Hyde et al. 2011; Ressler et al. 2022) and emerges from interactions among limbic affective drive, frontoparietal regulatory capacity, and subjective interpretations of past events (Ochsner and Gross 2005; Bishop 2007; Pessoa 2017). Understanding how these mechanisms jointly shape variation in emotional reactivity and vulnerability remains a central challenge in affective neuroscience.

Monoaminergic genes provide a window into these processes. Genetic variation affecting neurotransmitter storage or release can modulate affective reactivity (Pezawas et al. 2005; Suri et al. 2015). Among such genes, *SLC18A1* encodes vesicular monoamine transporter 1 (VMAT1), which transports monoamines into synaptic vesicles and thereby regulates the amplitude and timing of monoaminergic signaling (Erickson et al. 1996; Liu and Edwards 1997). Although VMAT2 is the dominant transporter in classical monoaminergic neurons, VMAT1 is also expressed in limbic and thalamic regions involved in emotional salience (Peter et al. 1995; Lohoff et al. 2006). Functional work shows that VMAT1 influences hippocampal neurogenesis and prefrontal dopaminergic signaling (Multani et al. 2013; Lohoff et al. 2019), suggesting broader involvement in central affective processes than previously assumed. A common nonsynonymous variant, Thr136Ile (rs1390938), alters transporter efficiency; the 136Ile allele increases monoamine uptake compared with 136Thr (Khalifa et al. 2012; Lohoff et al. 2014; Sato et al. 2019). This variant has also been associated with altered amygdala reactivity, functional connectivity, and white-matter microstructure (Lohoff et al. 2014; Zhu et al. 2015; Won et al. 2017), as well as the risk for affective, alcohol-use, bipolar, and autism spectrum disorders (Lohoff et al. 2006; Vaht et al. 2016; Noroozi et al. 2017). Moreover, evolutionary analyses suggest long-term maintenance of the polymorphism (Sato and Kawata 2018), possibly reflecting environmentally contingent phenotypic effects, consistent with evidence that emotional genetics often manifest most strongly in specific contexts.

Exposure to traumatic or stressful events is one of the major environmental determinants of emotional functioning. Yet psychological outcomes after adversity depend only weakly on exposure severity and more strongly on subjective cognitive–emotional processing, such as appraisal, perceived threat, and the vividness or intrusiveness of traumatic memory (Ehlers and Clark 2000; Brewin et al. 2010; Bonanno et al. 2011). These processes arise from amygdala–hippocampal–prefrontal mechanisms involved in emotional learning and autobiographical memory (Phelps 2004; Maren and Holmes 2016). Individuals differ substantially in how strongly these circuits engage during stressful experiences, leading some to encode adversity more vividly or interpret it as more threatening, which in turn contributes to variability in later emotional outcomes. Genetic factors that influence monoaminergic modulation may therefore shape emotional vulnerability by affecting the sensitivity of these neural systems to stress-related cues.

Neuroanatomical structure adds another source of variability. Cortical surface area and thickness differ across individuals due to developmental processes such as radial-unit proliferation and synaptic pruning (Rakic 1995; Petanjek et al. 2011). Regional differences in insular, parietal, and frontal cortices are linked to interoceptive awareness, salience detection, attentional control, and socioemotional integration (Seeley et al. 2007; Menon and Uddin 2010). Large-scale neuroimaging studies show that reduced cortical surface area or thickness is associated with greater vulnerability to depression, anxiety, and stress sensitivity (Goodkind et al. 2015; Schmaal et al. 2017; de Lange et al. 2025). These results support models in which emotional vulnerability reflects interactions between limbic affective reactivity and cortical regulatory capacity, with limited cortical resources amplifying emotional responses (Ochsner and Gross 2005; Pessoa 2017).

Progress in linking genetic variation, trauma processing, and cortical structure has been limited by the scarcity of datasets containing all three elements at population scale. The Tohoku Medical Megabank Project (TMM), established in the aftermath of the 2011 Great East Japan Earthquake (GEJE), was founded on a mission beyond purely academic inquiry. It was launched to monitor the long-term health and mental well-being of the disaster-affected population, aiming to inform individual- and community-level health initiatives and public health policies for recovery. Through this sustained commitment, the TMM has recruited tens of thousands of residents, creating a massive, integrated database that combines genomic data, neuroimaging, and extensive lifestyle questionnaires, including detailed items on earthquake-related disruption, evacuation, loss, and traumatic memory (Kuriyama et al. 2016; Hozawa et al. 2021). This unique endeavor to support community recovery has yielded a dataset of unprecedented depth and scale, which now serves as a vital resource for addressing the long-standing gap in our understanding of how biological and environmental factors interact. Unlike most cohorts that aggregate heterogeneous psychosocial stressors and life events, the TMM enables the psychological processing of a single, catastrophic event, the GEJE, to be quantified across both objective and subjective dimensions in an immense population. High-quality MRI measures of cortical morphology and comprehensive psychometric assessments further allow a multidimensional examination of how genetic variation, brain structure, and subjective interpretations of adversity shape emotional functioning. In this context, variation in monoaminergic genes such as *VMAT1* can be evaluated for both main effects and interactions with traumatic experiences and cortical regulatory systems.

The present study leverages the multimodal TMM cohort to examine how variation in the *VMAT1* rs1390938 polymorphism relates to affective functioning within a biopsychosocial framework. Our analyses were exploratory and aimed to clarify how variation in *VMAT1* relates to affective functioning through its combined effects with cortical morphology and trauma-related processes. By integrating genotype, MRI-derived cortical measures, psychological assessments, and detailed social and earthquake-related indices, we use the Thr136Ile variant as a biological probe to examine how genetic variation, brain structure, and lived experience jointly contribute to emotional diversity within a population. This multidimensional approach aims to illuminate the mechanisms through which variation in monoaminergic signaling becomes embedded in neurobiological and psychosocial pathways, giving rise to individual differences in reactivity, resilience, and vulnerability.

## Results

### Modest effect of *VMAT1* 136Thr on psychiatric outcomes

In the present cohort, the distribution of *VMAT1* genotypes yielded an approximately 3:1 ratio of the Thr to Ile alleles (Table S1). This pattern is consistent with allele frequencies previously reported in East Asian populations (Sato and Kawata 2018). To assess whether the association between *VMAT1* Thr136Ile and depressive symptoms reported in previous studies is recapitulated in our cohort, we first compared raw K6 scores between Ile carriers and Thr/Thr individuals (Fig. 1a). In females, K6 scores did not differ significantly between genotypes (Wilcoxon’s rank-sum test, *P* = 0.78). In males, however, Thr/Thr individuals showed significantly higher K6 scores than Ile carriers (*P* = 0.032), indicating a male-specific increase in depressive symptom severity associated with the 136Thr homozygous genotype.

**Figure 1.**
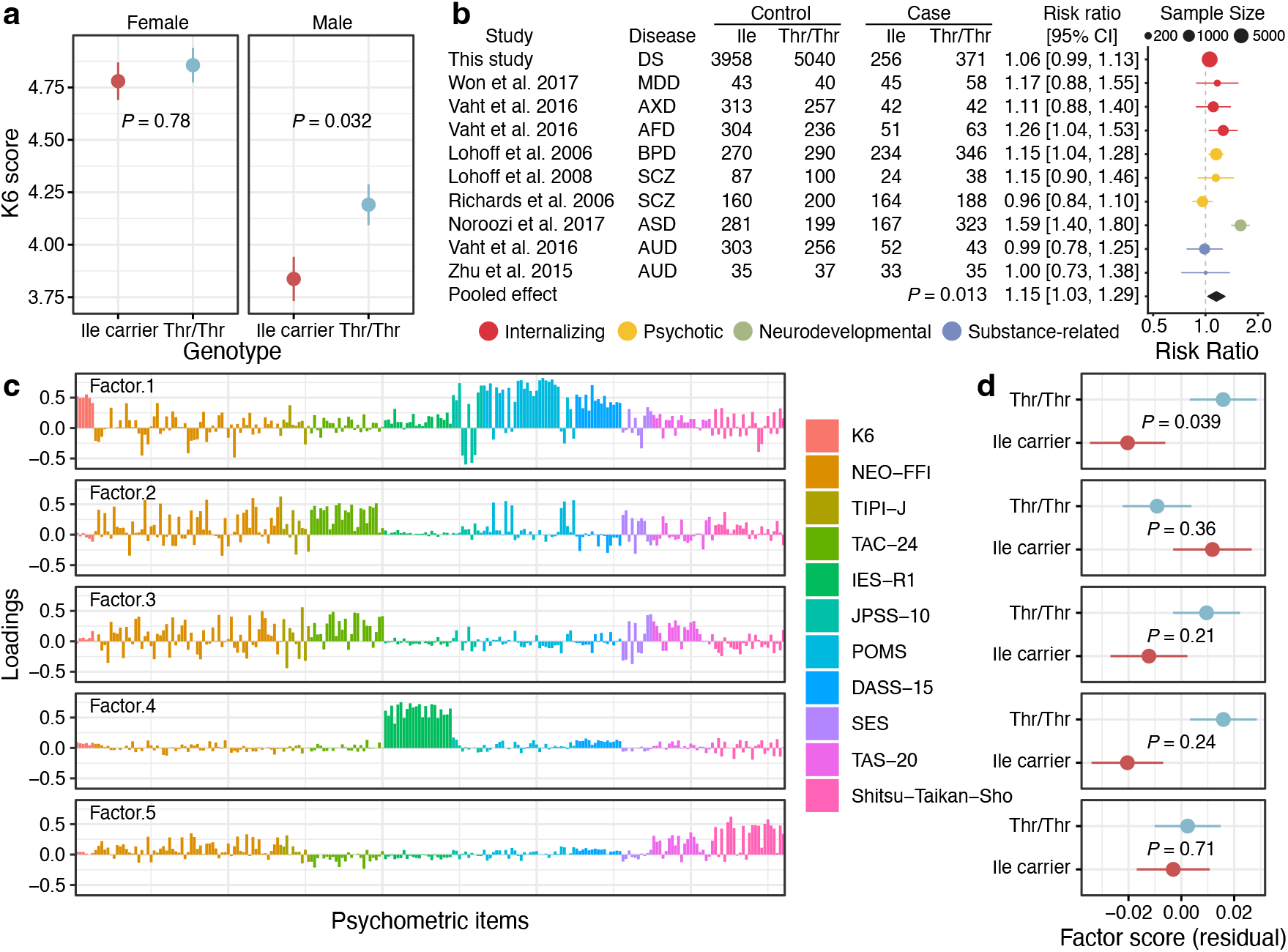
Associations between the *VMAT1* 136Thr/Thr genotype and affective vulnerability. **(a)** Sex‐stratified comparisons of K6 scores between Ile carriers and Thr/Thr individuals. Error bars represent the mean ± SE. **(b)** Meta-analysis of reported associations between the Thr136Ile variant and psychiatric conditions, including depressive state (DS), major depressive disorder (MDD), anxiety disorder (AXD), affective disorder (AFD), bipolar disorder (BPD), schizophrenia (SCZ), autism spectrum disorder (ASD), and alcohol use disorder (AUD). For the present cohort, cases were defined as individuals with K6 scores ≥ 13. Sample sizes for each genotype group (Ile carriers vs. Thr/Thr) are shown for controls and cases. Effect sizes are presented as risk ratios with 95% confidence intervals, with dot size indicating sample size. The pooled effect is shown at the bottom as a black diamond. **(c)** Factor loadings for the five latent factors extracted from multiple psychometric scales (K6, NEO-FFI, TIPI-J, TAC-24, IES-R1, JPSS-10, POMS, DASS-15, SES, TAS-20, Shitsu-Taikan-Sho). **(d)** Genotype effects on the five latent factor scores (corrected for sex and age), with *P*-values from Wilcoxon’s rank-sum tests.

We next evaluated the association between the *VMAT1* Thr136Ile variant and psychiatric case status by performing a meta-analysis that combined seven previously published studies (Lohoff et al. 2006; Richards et al. 2006; Lohoff et al. 2008; Zhu et al. 2015; Vaht et al. 2016; Noroozi et al. 2017; Won et al. 2017) with the present dataset assessing depressive state (K6 ≥ 13). Across all study arms, a random-effects meta-analysis indicated a significant overall association between the 136Thr/Thr genotype and increased risk of psychiatric case status (*P* = 0.013; Fig. 1b). To explore whether this association differed across diagnostic categories, we conducted group-specific meta-analyses without imposing a reference category. This analysis revealed a significant elevation of risk in study arms classified as Internalizing disorders (log(RR) = 0.107, 95% CI = [0.010, 0.203], *P* = 0.030) and an even stronger association in Neurodevelopmental disorders (log(RR) = 0.464, 95% CI = [0.340, 0.588], *P* < 2.2 × 10^−13^). In contrast, no significant associations were observed for Psychotic or Substance-related disorders, for which confidence intervals included zero. These findings suggest that the association between the *VMAT1* Thr136Ile variant and psychiatric outcomes may be more pronounced in diagnostic categories characterized by affective and developmental phenotypes. However, given the limited number of studies in some diagnostic groups and the presence of residual heterogeneity across study arms, these subgroup-specific effects should be interpreted cautiously and regarded as exploratory.

### Personality traits associated with the *VMAT1* 136Thr allele

Exploratory factor analysis of the psychometric battery identified five latent dimensions (Fig. 1c). Factor 1 primarily reflected negative affectivity and internalizing distress, with high loadings on K6, NEO-FFI Neuroticism, JPSS-10, POMS, DASS. Factor 2 captured interpersonal engagement and social positivity (NEO-FFI Extraversion, TIPI-J Items 1/5/6, TAC24). Factor 3 indexed self-regulation and behavioral control (NEO-FFI Agreeableness/Conscientiousness, TIPI-J Item 2/3/8, TAC24 items related to problem solving and emotional regulation). Factor 4 corresponded to trauma-related intrusion (IES-R1), and Factor 5 reflected somatic sensitivity (Shitsu-Taikan-Sho). The resulting factor scores were used for subsequent association tests with the *VMAT1* Thr136Ile genotype. To reduce potential confounding by demographic variables, factor scores were residualized for age and sex before association testing. When comparing these age- and sex-corrected factor scores across genotypes (Fig. 1d), individuals homozygous for the 136Thr allele showed higher scores on Factor 1, which was characterized by items reflecting negative affectivity and stress sensitivity. This genotype difference reached nominal significance (*P* = 0.039), although effects on the remaining four factors were not statistically significant (all *P* > 0.1). The direction of these associations is consistent with previous reports linking the *VMAT1* 136Thr allele to heightened emotional reactivity, suggesting that the variant may subtly modulate affective personality traits even within a non-clinical population.

### Cortical morphology associated with *VMAT1*-linked negative affectivity

Given that individuals with the *VMAT1* Thr/Thr genotype showed elevated scores on personality factor 1, which reflects negative affectivity and internalizing distress, we next examined whether variation in cortical surface area was associated with this psychological dimension. A whole-brain surface-to-factor 1 analysis in the Thr/Thr group revealed significant associations in three left-hemisphere regions: the fusiform, insula, and paracentral cortices (Benjamini-Hochberg *q* < 0.05; Fig. 2a). These regions showed the strongest negative associations, indicating that reduced surface area was linked to higher levels of negative affectivity in Thr/Thr individuals.

**Figure 2.**
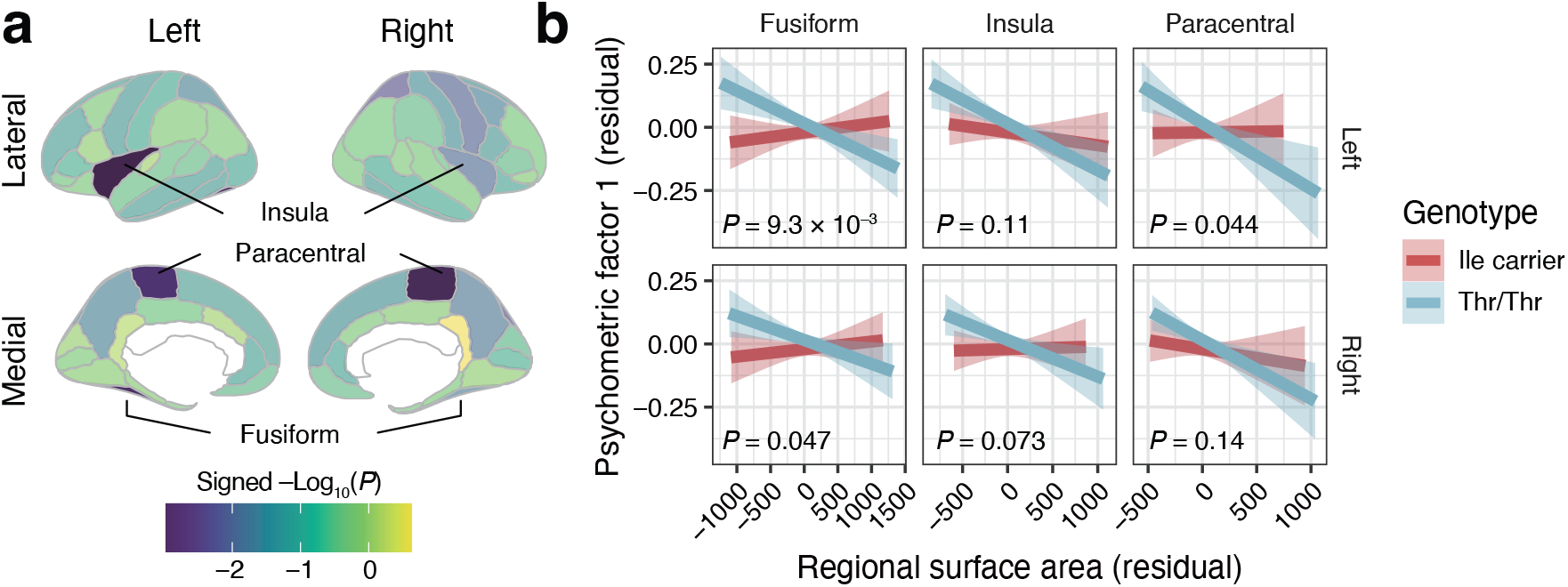
*VMAT1* genotype modulates the relationship between cortical surface area and psychometric factor 1. **(a)** Mapping of cortical regions showing significant associations between surface area and psychometric factor 1 scores of individuals with the Thr/Thr genotype. Color represents the signed –log_10_(*P*) values, indicating the direction and magnitude of association across lateral and medial views. Color opacity reflects significance based on Benjamini–Hochberg adjusted *q*-values, with semi-transparent regions denoting non-significant effects (*q* ≥ 0.05). **(b)** Genotype-dependent associations between regional surface area (fusiform, insula, paracentral; left and right hemispheres) and psychometric factor 1 scores, both corrected for age and sex. Shaded bands represent 95% confidence intervals (CIs). *P*-values are from linear interaction models testing genotype × surface area effects.

To determine whether these structure-to-trait relationships differed by genotype, we tested genotype-by-surface-area interaction effects for the same regions (Fig. 2b). Significant interactions were observed in the fusiform cortex in both hemispheres (left: *P* = 9.3 × 10^−3^; right: *P* = 0.047), as well as in the left paracentral cortex (*P* = 0.044). These results indicate that the relationship between cortical morphology and personality factor 1 varied depending on *VMAT1* Thr136Ile genotype. Specifically, Thr/Thr individuals showed steeper negative associations between surface area and negative affectivity compared with Ile carriers. Taken together, these findings suggest that structural variation in the fusiform and paracentral cortices moderates the behavioral impact of *VMAT1* Thr136Ile, linking genetic variation to individual differences in emotional reactivity.

Importantly, this moderation did not arise from baseline morphological differences between genotype groups. Exploratory whole-brain comparisons revealed no cortical region in which surface area significantly differed between *VMAT1* genotypes in a Wilcoxon rank-sum test (Fig. S1), indicating that the genotype did not exert broad main effects on cortical morphology. Rather, *VMAT1* genotype influenced how inter-individual variation in cortical structure related to negative affectivity, consistent with a context-dependent rather than a global structural effect.

### Cortical morphology modulates *VMAT1*-by-trauma effects on depressive symptoms

To identify which environmental components interact with *VMAT1* genotype to influence depressive symptoms, we first tested all four principal components derived from the questionnaire datasets (two PCs from social-relationship items and two PCs from GEJE-related items). Social-relationship PC1, where higher scores indicate stronger social connectedness and more supportive interpersonal relationships, reflected overall social embeddedness. Social-relationship PC2 captured variation in household composition, with higher scores corresponding to a greater number of cohabiting family members and more restricted non-cohabiting social ties (Fig. S2a). GEJE-related PC1 indexed the overall severity of disaster-related disruption, such that higher PC1 values reflected less objective damage or adversity experienced during the GEJE. GEJE-related PC2 represented subjective trauma sensitivity, defined as the discrepancy between objective exposure and current traumatic memory; higher PC2 scores corresponded to weaker or less persistent traumatic memory despite the level of objective exposure (Figs. 3a and S2b).

**Figure 3.**
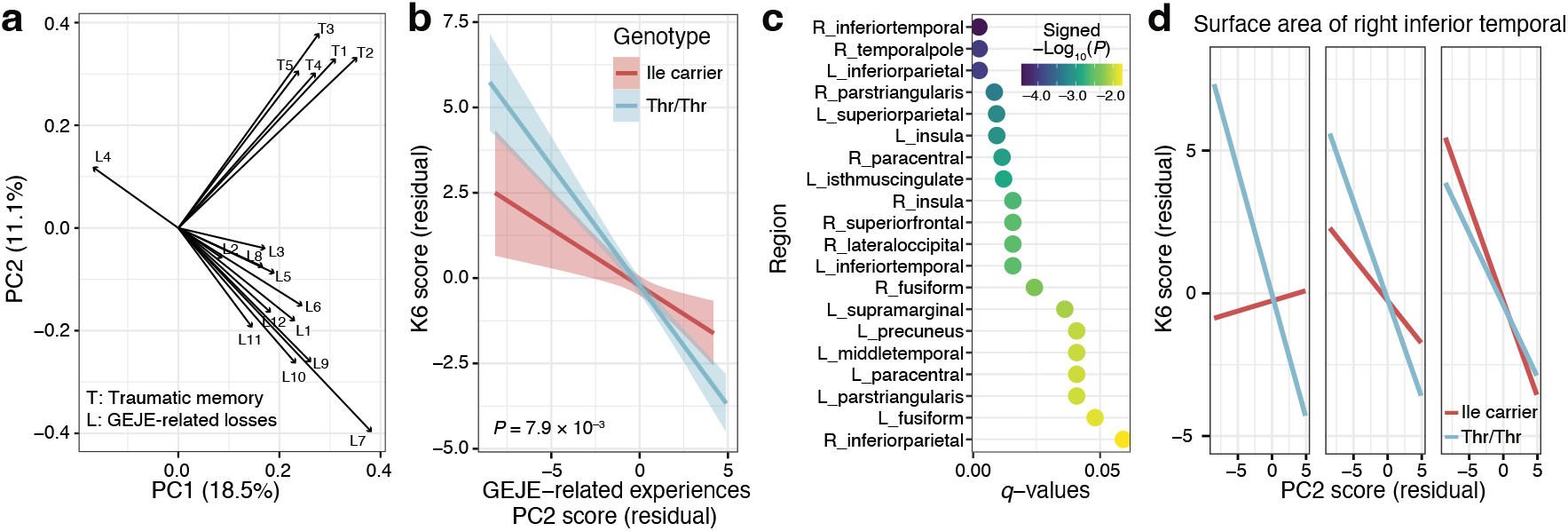
Subjective trauma sensitivity, *VMAT1* genotype, and cortical surface area jointly shape depressive symptoms. **(a)** Principal component analysis (PCA) of questionnaires regarding GEJE-related experience. L-variables represent objective disaster-related exposures and losses associated with the Great East Japan Earthquake (GEJE), whereas T-variables represent current traumatic memory. PC1 captures the severity of GEJE-related experiences, whereas PC2 reflects the relative magnitude of subjective traumatic memory compared with the objective severity of disaster exposure. **(b)** Relationship between age- and sex-corrected PC2 and K6 scores across genotypes. The *P*-value reflects the PC2 × genotype (Ile carrier vs. Thr/Thr) interaction term, indicating genotype-dependent effects of PC2 on K6. Shaded bands represent 95% CIs. **(c)** Cortical regions in which the three-way interaction between genotype, PC2, and regional surface area significantly predicts depressive symptoms. Color represents signed –log_10_(*P*) values, highlighting regions where surface area moderates the genotype-dependent effect of subjective trauma sensitivity on depression. Region labels are prefixed with L_ and R_ to denote the left and right hemispheres, respectively. **(d)** Predicted K6 scores plotted against PC2 values at –1 SD, mean, and +1 SD of right inferior temporal surface area, illustrating genotype-dependent modulation of the PC2–depression relationship.

Before conducting the interaction analyses, we examined the broader associations between these PCs and psychological traits. Social-relationship PC1 showed strong correlations with personality Factors 1–3 (*r* = –0.28, 0.39, –0.26, respectively; all *P* < 0.001; Fig. S2c), whereas social-relationship PC2 showed only weak associations with any psychological factor, with absolute *r* values of approximately 0.1. GEJE-related PC1 was associated with negative affectivity corresponding to Factor 1 (*r* = –0.20, *P* < 0.001) and, more strongly, with PTSD-related symptoms corresponding to Factor 4 (*r* = –0.48, *P* < 0.001). GEJE-related PC2 was also correlated with Factor 4 (*r* = –0.19, *P* < 0.001), although the association was weaker than that of PC1. None of the PCs differed significantly by *VMAT1* genotype (Fig. S2d). Thus, the environmental dimensions themselves were largely independent of genotype, providing a clean context for evaluating gene-by-environment effects.

Each PC score was then tested for a Genotype × PC interaction effect on age- and sex-residualized K6 depressive symptoms. Among the four components, only the GEJE-related PC2 showed a significant interaction with *VMAT1* genotype (Fig. 3b). Individuals with the Thr/Thr genotype exhibited a steeper positive relationship between PC2 and K6 scores compared with Ile carriers, and this genotype-dependent effect was particularly pronounced at lower PC2 values, where subjective trauma was stronger. No interaction effects were observed for the social components (PC1 or PC2) or for the GEJE-related PC1 (all *P* > 0.1; Fig. S3). Because GEJE-related PC2 was the only dimension showing a robust genotype-dependent effect on depressive symptoms, we focused subsequent analyses on whether cortical morphology further moderated this PC2 × genotype interaction. A whole-brain screening of genotype-by-PC2-by-surface-area interactions identified several regions in which this three-way interaction significantly predicted depressive symptoms (Fig. 3c). Notably, right inferior temporal lobule showed the strongest effect after Benjamini-Hochberg correction, with a corresponding effect also observed in the left inferior temporal lobule.

Additional interactions were observed bilaterally in the insula, fusiform gyrus, paracentral cortex, pars triangularis, and inferior parietal lobule. In all regions with significant or suggestive effects, the interaction term was negative. This pattern indicates that the genotype-dependent effect of subjective trauma sensitivity on depressive symptoms was amplified when regional surface area was smaller and attenuated as surface area increased. As shown in Fig. 3d, Thr/Thr individuals exhibited a steeper positive association between PC2 and depressive symptoms at lower levels of inferior temporal surface area, whereas this genotypic difference was minimal at larger surface areas. These findings suggest that regional cortical morphology moderates the behavioral impact of traumatic experiences in a genotype-dependent manner.

Finally, to clarify whether *VMAT1* genotype and cortical structure exerted broad interactive effects on PC scores themselves, we tested genotype-by-surface-area effects on all PC scores. No significant interactions were observed after multiple correction (Fig. S2e), indicating that *VMAT1* genotype did not influence psychosocial and environmental dimensions either directly or through its interaction with cortical morphology. Together, these findings suggest that regional cortical morphology moderates the behavioral impact of traumatic experiences in a genotype-dependent manner, rather than shaping the environmentally derived components themselves.

## Discussion

The present study demonstrates that genetic variation in *VMAT1*, specifically the Thr136Ile polymorphism, contributes to individual differences in affective functioning through its interplay with cortical morphology and sensitivity to traumatic experiences. Using a large population-based cohort enriched with multimodal data from the Tohoku Medical Megabank Project, we show that the 136Thr homozygous genotype is associated with higher levels of depressive symptoms and negative affectivity, consistent with previous reports linking this allele to heightened emotional reactivity and psychiatric vulnerability (Lohoff et al. 2006; Richards et al. 2006; Lohoff et al. 2014; Vaht et al. 2016; Noroozi et al. 2017). Notably, the association between *VMAT1* genotype and depressive tendency was observed only in males; however, sex-specific effects of VMAT1 have not been consistently reported in previous studies, and it remains unclear whether this pattern reflects population-specific factors in the Japanese cohort or more general sex-dependent mechanisms, warranting further investigation. Our analysis also provides new evidence that the emotional effects of this genetic variant are significantly amplified by structural features of the cortex and differences in subjective trauma-related processing.

Specifically, our findings reveal that emotional vulnerability is not determined by any single factor, but rather arises from interactions among three distinct layers: genetic background, neurobiological architecture, and subjective processing of life experiences. Factor analysis of psychometric measures indicated that personality factor 1, a latent dimension reflecting negative affectivity and internalizing distress, differed by *VMAT1* genotype, with 136Thr/Thr individuals exhibiting higher scores. This pattern is consistent with prior reports linking the Thr allele to heightened emotional reactivity and stress-sensitive temperament (Lohoff et al. 2014; Vaht et al. 2016). At the same time, our imaging and interaction analyses demonstrate that this genetic predisposition does not act in isolation. Instead, regional cortical morphology appears to function as a neurobiological buffer against emotional distress. Our imaging analyses further showed that, within the Thr/Thr group, smaller surface area in the fusiform, insula, and paracentral cortices was associated with higher negative affectivity, suggesting that regional cortical morphology moderates the expression of VMAT1-related affective tendencies. Crucially, this means that even individuals with high genetic risk may remain resilient if they possess sufficient cortical structural capacity to integrate and regulate affective signals. Conversely, limited cortical resources act as a constraint, amplifying the behavioral impact of genetic and environmental risks.

Although VMAT1 expression is most consistently documented in limbic and subcortical structures, including the amygdala, hippocampus, and thalamus (Peter et al. 1995; Lohoff et al. 2006), the cortical pattern observed here can be interpreted within broader corticolimbic network organization. The amygdala, a key locus of emotional salience detection (LeDoux 2000; Phelps and LeDoux 2005), is sensitive to monoaminergic modulation (Pezawas et al. 2005; Munafò et al. 2008; Liu et al. 2018), and the Thr136Ile genotype has been shown to influence its activity (Lohoff et al. 2014; Sato et al. 2022). The insula, which maintains strong structural and functional connectivity with the amygdala (Baur et al. 2013; Denny et al. 2014), can transmit limbic signals to higher-order cortical systems involved in appraisal and regulatory control (Critchley et al. 2004; Paulus and Stein 2006; Kober et al. 2008; Craig 2009). The fusiform cortex supports the processing of socially salient facial information (Kanwisher and Yovel 2006; Jung et al. 2021), while the paracentral cortex participates in stress-related integration within midline and parietal networks (Buhle et al. 2014; Zhang et al. 2020; Tang et al. 2024). The emergence of VMAT1-related effects in these regions suggests that monoaminergic variation may shape emotional functioning through the distributed cortical systems that interpret and regulate limbic signals.

This pattern raises the possibility that cortical surface area indexes aspects of network capacity relevant for integrating and contextualizing affective information. Cortical surface area is thought to reflect a higher number of ontogenetic columns (Rakic 1995), which may partly translate to increased processing capacity within large-scale cortical networks (Murray et al. 2024). Larger surface area in regions such as the insula or fusiform cortex may therefore support more effective multimodal integration, reappraisal, and contextualization of affective cues, whereas smaller surface area may constrain these operations. Under this interpretation, *VMAT1* genotype may contribute to the intensity or reactivity of limbic affective signals, while cortical morphology determines the regulatory bandwidth available to modulate those signals. The interaction effects observed here, in which smaller regional surface area amplified VMAT1-related emotional tendencies, are consistent with multiplicative models in which emotional vulnerability arises from heightened bottom-up reactivity combined with limited top-down integrative resources (Ochsner and Gross 2005; Bishop 2007; Pessoa 2017; LeDuke et al. 2023).

The role of GEJE-related PC2 adds a further, independent dimension to this multilevel architecture. PC2 captures the discrepancy between objective disruption and the strength of current traumatic memory, indexing subjective trauma encoding rather than exposure severity. Cognitive models of trauma describe how disproportionate or overgeneralized traumatic memory can arise from heightened emotional reactivity or maladaptive appraisal processes (Ehlers and Clark 2000; Bonanno et al. 2011). In the present study, this dimension interacted with *VMAT1* genotype and was further moderated by cortical morphology in a distributed set of regions. The inferior temporal and parietal cortices, together with the insula and fusiform regions, form part of a broader network supporting semantic representation, self-referential evaluation, attentional control, and multimodal integration (Corbetta et al. 2008; Kober et al. 2008; Pessoa 2017). These functions are well suited for shaping the interpretation and contextualization of autobiographical memory, including traumatic experiences. Within this framework, individuals with heightened monoaminergic reactivity (Thr/Thr genotype), stronger subjective trauma encoding (low PC2 scores), and reduced cortical network capacity in these regions may experience disproportionate psychological impact even when objective adversity is relatively modest. This pattern illustrates how genetic variation, subjective experience, and distributed cortical structure jointly amplify vulnerability in ways that would not be detectable when examined in isolation.

Finally, it is worth considering how these findings relate to the evolutionary history of *VMAT1* and other monoaminergic genes. In the present study, the average effect of the Thr136Ile variant on psychological traits was modest, yet its influence became more apparent in specific environmental and neurobiological contexts. This pattern is consistent with the fact that recent large-scale GWAS of personality traits have not identified significant associations at this locus, as well as several historically prominent monoaminergic candidate genes, such as *SLC6A4, MAOA*, and *COMT* (Lo et al. 2017; Nagel et al. 2018; Gupta et al. 2024; Schwaba, Clapp Sullivan, et al. 2025). Contemporary GWAS typically estimate effect sizes within a particular cultural or environmental background or across heterogeneous populations, which can obscure the influence of genes whose effects are strongly context dependent. Moreover, since personality and affective traits are highly polygenic (Zeng et al. 2021; Zietsch 2024), the contribution of any single locus tends to be small when averaged across diverse environmental settings. This polygenic architecture further makes it difficult to detect context-dependent genetic effects in large-scale GWAS, helping to explain why monoaminergic candidate genes often show limited signal in such studies despite exhibiting more pronounced effects under specific environmental and neurobiological conditions. At the same time, candidate-gene approaches in psychiatric genetics have a mixed track record, and our findings should be interpreted as complementary to, rather than in opposition to, insights from genome-wide studies and ideally be followed by tests in polygenic frameworks. These context-dependent genetic effects are also compatible with the evolutionary history of *VMAT1*. In ancestral environments in which ecological and social conditions were more uniform, such as during early phases of human evolution or the initial out-of-Africa expansion, the context-dependent value of affective tendencies may have been more consistent, potentially making certain behavioral profiles relatively advantageous (Sato and Kawata 2018; Sato et al. 2019). In contrast, modern environments exhibit substantial spatial and temporal heterogeneity, making the adaptive value of emotional and behavioral tendencies more situational. Such variability in selective contexts may contribute to the long-term maintenance of genetic diversity at loci like *VMAT1* (Sato and Kawata 2018), even when their average effect sizes in contemporary populations appear small.

Despite these significant advances, several limitations warrant consideration. First, the analyses were cross-sectional, which prevents causal inference about the relationships among *VMAT1* genotype, cortical structure, and trauma-related psychological outcomes. Second, while genomic and MRI data were available for most participants in the cohort, detailed lifestyle questionnaire data, including items related to earthquake exposure and traumatic memory, were available for a smaller subset. The limited sample size for trauma-related measures may reduce the statistical power of the gene-by-environment analyses. Third, the cortical findings were derived from whole-brain exploratory screenings, and although we controlled for multiple comparisons using false discovery rate procedures, the number of statistical tests was large, and the results should be viewed as hypothesis-generating and in need of replication. Finally, because the TMM cohort was established in the aftermath of the 2011 Great East Japan Earthquake, shared environmental context may influence the psychological and neural characteristics of the sample. Future longitudinal and cross-cultural studies will be necessary to evaluate the extent to which the present findings extend beyond this population.

Taken together, the present findings outline a multilevel and multiplicative pathway through which variation in *VMAT1* may contribute to emotional vulnerability. The 136Thr allele was associated with heightened negative affectivity and depressive symptoms, and these tendencies were shaped by both subjective trauma processing and cortical morphology in regions supporting emotional appraisal and memory integration. This framework highlights how genetic variation interacts with individual interpretations of past experiences as well with neurobiological substrates to jointly shape affective responses to adversity. By integrating genomic, psychological, and neuroimaging data from a large population-based cohort, this study underscores the value of multimodal approaches for understanding risk and resilience and provides a foundation for future research aimed at clarifying how biological predispositions and experiential factors jointly shape variation in mental health.

## Materials and methods

### A large-scale cohort dataset of Tohoku Medical Megabank Organization

In this study, we used genome array, neuroimaging, and lifestyle questionnaire data from participants of the Tohoku Medical Megabank Project (TMM) conducted by the Tohoku Medical Megabank Organization. The dataset and sample extraction flow are summarized in Fig. S4a. Genome data and genotype of the *VMAT1* polymorphism of interest in this study (Thr136Ile; rs1390938) were obtained from the Japonica Array v2 and v3, population-optimized SNP arrays enriched for functional variants in the Japanese population. To enable sample-level quality control based on genome-wide markers, all genotyped individuals were merged with reference samples from the 1000 Genomes Project reference panel (1000 Genomes Project Consortium et al. 2015), and quality control for SNPs was performed using PLINK2 (Chang et al. 2015). Variants with a minor allele frequency below 0.05, a genotype missingness rate above 2%, or deviation from Hardy–Weinberg equilibrium (*P* < 1 × 10^−6^) were excluded. Linkage disequilibrium pruning was conducted using a 200-kb window, 50-variant step size, and *r*^2^ threshold of 0.2. For the remaining 115,893 SNPs, principal component analysis (PCA) was performed to identify and exclude participants whose genetic background deviated from the cluster representing the Japanese population (Fig. S4b). We also excluded monozygotic-twin–like or duplicate samples using KING-robust relatedness inference (--king-cutoff 0.353) as indicated (Manichaikul et al. 2010).

After these quality control procedures, we utilized the MRI 12K dataset, which includes high-resolution brain magnetic resonance imaging and cognitive assessment data. Participants with missing values in either the MRI or cognitive variables were excluded, resulting in a total of 9,625 individuals (6,046 females and 3,579 males) available for the main analyses. For a subset of these participants, we further obtained lifestyle questionnaire data from two major TMM cohort studies: the TMM Community-Based Cohort Study (TMM CommCohort Study) and the TMM Birth and Three-Generation Cohort Study (TMM BirThree Cohort Study). From these datasets, we extracted information related to social relationships (including cohabiting family members) and earthquake experiences (including subjective memories of the Great East Japan Earthquake). After excluding individuals with missing responses in any of these variables, a total of 2,229 participants (1,496 females and 733 males) were included in the analyses involving lifestyle data. The number of individuals in each genotype group is summarized in Table S1, showing an approximate 3:1 ratio of Thr to Ile alleles, which is overall consistent with allele frequencies reported in East Asian populations (Sato and Kawata 2018).

### General lifestyle questionnaire dataset

Participants completed a self‐administered lifestyle questionnaire that was part of the baseline assessment of the TMM cohort studies. The social-relationship section assessed aspects such as co-residence patterns, frequency and quality of interactions with family, relatives, and friends, perceived availability of emotional or practical support, motivation to form new social ties, and perceptions of neighborhood cohesion. Items assessing the frequency of contact, emotional closeness, and perceived support from relatives and friends corresponded to the Lubben Social Network Scale (LSNS-6) (Lubben et al. 2006; Kurimoto et al. 2011). Perceptions of neighborhood cohesion were assessed using items capturing mutual aid, trust, everyday social interaction, and collective problem solving within the community. The GEJE-related section assessed both objective and subjective dimensions of disaster exposure, including housing damage, post-disaster residential transitions, repeated relocations, exposure to damaged surroundings, direct or witnessed life-threatening events, bereavement due to the disaster, as well as subjective psychological responses such as intrusive memories, emotional distress, physiological reactivity, and avoidance behaviors related to the event (Ishikuro et al. 2022; Kotozaki et al. 2023). Responses were recorded using binary, categorical, or Likert-type formats, as appropriate. The questionnaire had been administered by TMM staff in cooperation with municipal health and research centers, with informed consent obtained at the time of enrollment.

To reduce dimensionality and uncover latent psychosocial patterns, we conducted PCA separately for social-relationship items and GEJE-related items. Prior to PCA, questionnaire responses were numerically coded according to the original response formats, with higher values reflecting either greater or lesser exposure, endorsement, or frequency depending on item wording. For binary items coded as yes = 1 and no = 2, positive loadings indicate the absence of the queried experience, whereas negative loadings indicate its presence. For social items, two components emerged: one reflecting overall social connectedness and another representing variation in household structure and the breadth of non-cohabiting social ties (Fig. S2a). For GEJE-related items, two components were extracted: one indexing the objective severity of GEJE exposure and the other capturing subjective trauma sensitivity, defined as the extent to which current traumatic memory diverged from objective exposure (Figs. 3a and S3b). All component scores were residualized for age and sex and used in the genotype-by-environment analyses described below.

### Psychometric dataset

To assess psychological traits and states relevant to our study, we administered a battery of standardized self-report questionnaires in Japanese. Psychological distress was measured using the Kessler Psychological Distress Scale (K6), a widely used 6-item instrument designed to assess nonspecific symptoms of mental distress (Kessler et al. 2002). Personality traits were assessed with the 60-item NEO Five-Factor Inventory (NEO-FFI), which evaluates the Big Five dimensions of Neuroticism, Extraversion, Openness to Experience, Agreeableness, and Conscientiousness (Costa and McCrae 1989), as well as the Japanese version of the Ten-Item Personality Inventory (TIPI-J), a brief instrument capturing the same five-factor structure (Gosling et al. 2003; Oshio et al. 2012). To evaluate coping strategies, we used the Tri-Axial Coping Scale (TAC-24), which was developed in Japan to assess problem-focused, emotion-focused, and avoidance-oriented coping tendencies (Takamoto and Aikawa 2012). Posttraumatic stress symptoms were assessed using the Japanese version of the Impact of Event Scale-Revised (IES-R), which includes items reflecting intrusion, avoidance, and hyperarousal (Weiss and Marmar 1997). Perceived stress was measured using the Japanese Perceived Stress Scale (JPSS-10), based on the original scale (Cohen et al. 1983). Mood states were evaluated using the Japanese version of the Profile of Mood States (POMS), covering tension, depression, anger, fatigue, confusion, and vigor subscales (Mcnair et al. 1971). Depression, anxiety, and stress symptoms were assessed using the 15-item short form of the Depression Anxiety Stress Scales (DASS-15), derived from the original 42-item version (Lovibond and Lovibond 1995). Global self-esteem was measured using the Rosenberg Self-Esteem Scale (SES (Rosenberg 1965)), and emotional awareness and alexithymia were assessed using the 20-item Toronto Alexithymia Scale (TAS-20) (Bagby et al. 1994). We also included the Somatic Awareness and Resilience Scale (Shitsu-Taikan-Sho), a 23-item Japanese-developed instrument assessing bodily awareness, self-regulation, and behavioral tendencies related to stress resilience (Arimura et al. 2012). All instruments were administered in Japanese, using validated translations when available, and responses were scored according to the original developers’ instructions.

To reduce the dimensionality of the psychometric dataset and to derive latent dimensions of personality and affective traits, we conducted exploratory factor analysis (EFA) on the standardized questionnaire scores. The analysis was performed using the fa() function in the *psych* package in R, with maximum likelihood estimation (fm = “ml”) and promax rotation (rotate = “promax”). The number of factors was determined to be 5, potentially corresponding to the five major personality dimensions. Individual factor scores were then computed for each participant and used as dependent variables in subsequent genotype–phenotype association analyses.

### Meta-analysis of the effect of *VMAT1* 136Thr on psychiatric disorders

We conducted a random-effects meta-analysis to examine the association between the Thr136Ile variant in *VMAT1* and psychiatric case–control status across previously published studies (Lohoff et al. 2006; Richards et al. 2006; Lohoff et al. 2008; Zhu et al. 2015; Vaht et al. 2016; Noroozi et al. 2017; Won et al. 2017), together with the present dataset. In our study, the depressive state of participants was screened using the K6 score, and individuals scoring 13 or higher were classified as cases, following established thresholds for serious psychological distress (Kessler et al. 2003). Genotypes were binarized such that individuals carrying at least one Ile allele (i.e., Ile/Ile or Thr/Ile) were grouped together and contrasted against those with the Thr/Thr genotype. This dichotomization was applied consistently across all included studies to enable comparable effect size estimation. Using genotype counts from case and control groups, log risk ratios (log(RR)) and corresponding sampling variances were computed with the escalc() function in the *metafor* package in R. To evaluate the overall association between the *VMAT1* Thr136Ile variant and psychiatric case status, we first fitted a random-effects meta-analysis model across all study arms. To explore potential differences across diagnostic categories, we then conducted exploratory subgroup-specific meta-analyses stratified by diagnostic group. Diagnostic categories were defined as Internalizing disorders (including depressive state, major depressive disorder, affective disorder, and anxiety disorder), Neurodevelopmental disorders (including autism spectrum disorder), Psychotic disorders (including schizophrenia and bipolar disorder), and Substance-related disorders (including alcohol use disorder), based on recent transdiagnostic work quantifying genome-wide genetic correlations among psychiatric disorders (Grotzinger et al. 2025; Schwaba, Mallard, et al. 2025). Within each diagnostic group, effect sizes were pooled using random-effects models without imposing a reference category. Between-study heterogeneity was assessed using Cochran’s *Q* statistic, *τ*^2^, and *I*^2^. All effect sizes are reported as log risk ratios comparing the 136Thr/Thr genotype with Ile-allele carriers.

### Cortical surface area analysis

We examined the association between cortical morphology and personality traits separately within each *VMAT1* genotype group. Regional cortical surface areas (68 Desikan–Killiany atlas regions) were extracted using FreeSurfer v7.2.0 after standard preprocessing, including intensity normalization, skull stripping, and white-matter and pial surface reconstruction. MRI data were acquired on the same day as the psychometric assessments. All regional surface-area measures were adjusted for total cortical surface area by linear regression, and residuals were used in subsequent analyses. First, to examine whether *VMAT1* genotype was associated with baseline variation in cortical morphology, adjusted surface-area values were compared between genotype groups (Ile carriers vs. Thr/Thr) using Wilcoxon rank-sum tests. Next, for the Thr/Thr group, partial correlation analyses were performed between adjusted regional surface area and personality factor 1 scores, controlling for age and sex. Multiple-comparison correction across regions was applied using the Benjamini–Hochberg false discovery rate (*q* < 0.05). To assess whether structure-to-trait relationships differed by genotype, we fitted linear interaction models of the form:

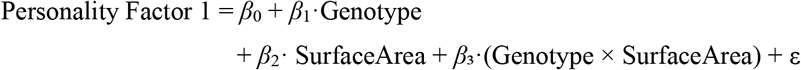

where Genotype was coded as 0 for Thr/Thr and 1 for Ile-allele carriers. The interaction term (*β*_3_) tested whether the slope relating cortical surface area to factor 1 scores differed between genotype groups.

### Exploratory analyses of lifestyle–experience components

To characterize how lifestyle–experience components relate to psychological characteristics, Pearson correlations were computed between each PC score and the five psychological factors derived from the psychometric battery. To test whether lifestyle–experience components differed by *VMAT1* genotype, PC scores were compared between Thr/Thr individuals and Ile carriers using Wilcoxon rank-sum tests. To evaluate whether cortical morphology and genotype jointly contributed to variation in lifestyle–experience components themselves, each PC score was regressed on genotype, regional surface area, and their interaction according to the following model:

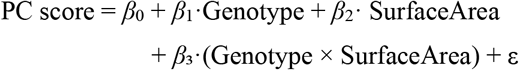

These exploratory analyses are summarized in Fig. S2.

### Gene-by-environment-by-brain interaction analysis

Given that GEJE-related PC2 was the only environmentally derived component showing a significant interaction with *VMAT1* genotype on depressive symptoms, we next examined whether cortical morphology further moderated this gene–environment effect. K6 scores, PC2 scores, and regional cortical surface-area measures were all residualized for age and sex before analysis. For each cortical region, we fitted the following linear model:

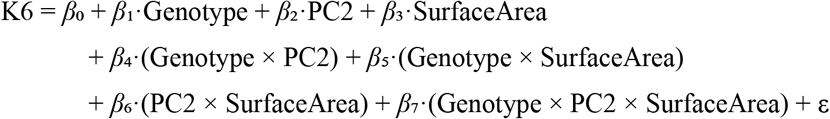

where Genotype was coded as 0 for Thr/Thr and 1 for Ile-allele carriers. The three-way interaction term (*β*_7_) tested whether the association between subjective trauma sensitivity (PC2) and depressive symptoms differed by genotype depending on regional cortical surface area. For each region, *P*-values for the three-way interaction were adjusted for multiple comparisons using the Benjamini–Hochberg procedure. The right inferior temporal lobule, which showed the strongest three-way interaction effect, was visualized using simple-slope plots across representative levels of surface area (mean ± 1 SD) separately for each genotype group.

### Statistical analysis

All statistical analyses were performed in R v4.4.2. Meta-analytic models were fitted using the *metafor* package. Random-effects models estimated log risk ratios, and heterogeneity was assessed using *Q, τ*^2^, and *I*^2^. Factor analyses were conducted using the *psych* package with maximum-likelihood estimation and promax rotation. Principal component analyses of the questionnaire items were performed on standardized variables, and PC scores were residualized for age and sex. Group differences were tested using Wilcoxon’s rank-sum tests. Associations between cortical surface area and personality factor scores were examined using partial correlations controlling for age and sex, followed by linear interaction models testing genotype-by-surface-area effects. Whole-brain multiple comparisons were corrected using the Benjamini–Hochberg false discovery rate. Three-way interaction models tested whether cortical morphology moderated the genotype-by-PC2 effect on depressive symptoms. For each region, significance of the Genotype × PC2 × SurfaceArea term was evaluated and FDR-corrected across regions. Effect visualization used simple-slope plots at mean ± 1 SD of surface area.

### Ethical statement

This study was conducted using data obtained from the Tohoku Medical Megabank Project (TMM), which was approved by the Ethics Committee of the Tohoku Medical Megabank Organization. The use of TMM data for the present study was also approved by the Ethics Committee of Tohoku University (Approval IDs: 2016-4-058 and 2020-1011). All participants provided written informed consent for participation in the TMM project, including the secondary use of their data for research purposes. The study complied with the principles of the Declaration of Helsinki.

## Acknowledgements

We thank the Tohoku Medical Megabank Organization and the participants donating their samples and information. This work was supported by a Grant-in-Aid for Scientific Research (17H05934 and 19H04892 to M.K.) by the Japan Society for the Promotion of Science.

## Author contributions

D.X.S. and M.K. conceived and designed the study. M.K. acquired the funding. D.X.S. conducted the data analysis and wrote the initial draft of the manuscript. T.M. and K.Ki. supervised genomic analyses; S.M., neuroimaging analyses; and Z.Y. and H.T., psychological data interpretation. K.Ka. contributed to the analysis of psychological data together with D.X.S. All authors reviewed and approved the final manuscript.

## Data and code availability

The individual data are under restricted access. Access can be obtained upon request after approval the Ethical Committee and the Materials and Information Distribution Review Committee of ToMMo. For detailed guidance on making the data access request, see website (http://www.dist.megabank.tohoku.ac.jp). Codes to reproduce the findings described in the present study will be available upon publication.

