## Supplementary Information for "Monoaminergic variation, cortical structure, and disaster trauma interact to shape emotional vulnerability"

This file includes:

- **Table S1**
- **Figures S1–4**

**Table S1. Demographic characteristics and *VMAT1* genotype distribution of the study sample.**

| | | Genotype frequencies | | | Allele frequencies | | Total |
|---|---|---|---|---|---|---|---|
| Dataset | Sex | Thr/Thr | Thr/Ile | Ile/Ile | Thr | Ile | |
| Dataset 1 | Male | 2,052 (57.3%) | 1,294 (36.2%) | 233 (6.5%) | 75.4% | 24.6% | 3,579 |
| | Female | 3,359 (55.6%) | 2,299 (38.0%) | 388 (6.4%) | 74.6% | 25.4% | 6,046 |
| Dataset 2 | Male | 411 (56.1%) | 279 (38.1%) | 43 (5.9%) | 75.1% | 24.9% | 733 |
| | Female | 835 (55.8%) | 570 (38.1%) | 91 (6.1%) | 74.9% | 25.1% | 1,496 |

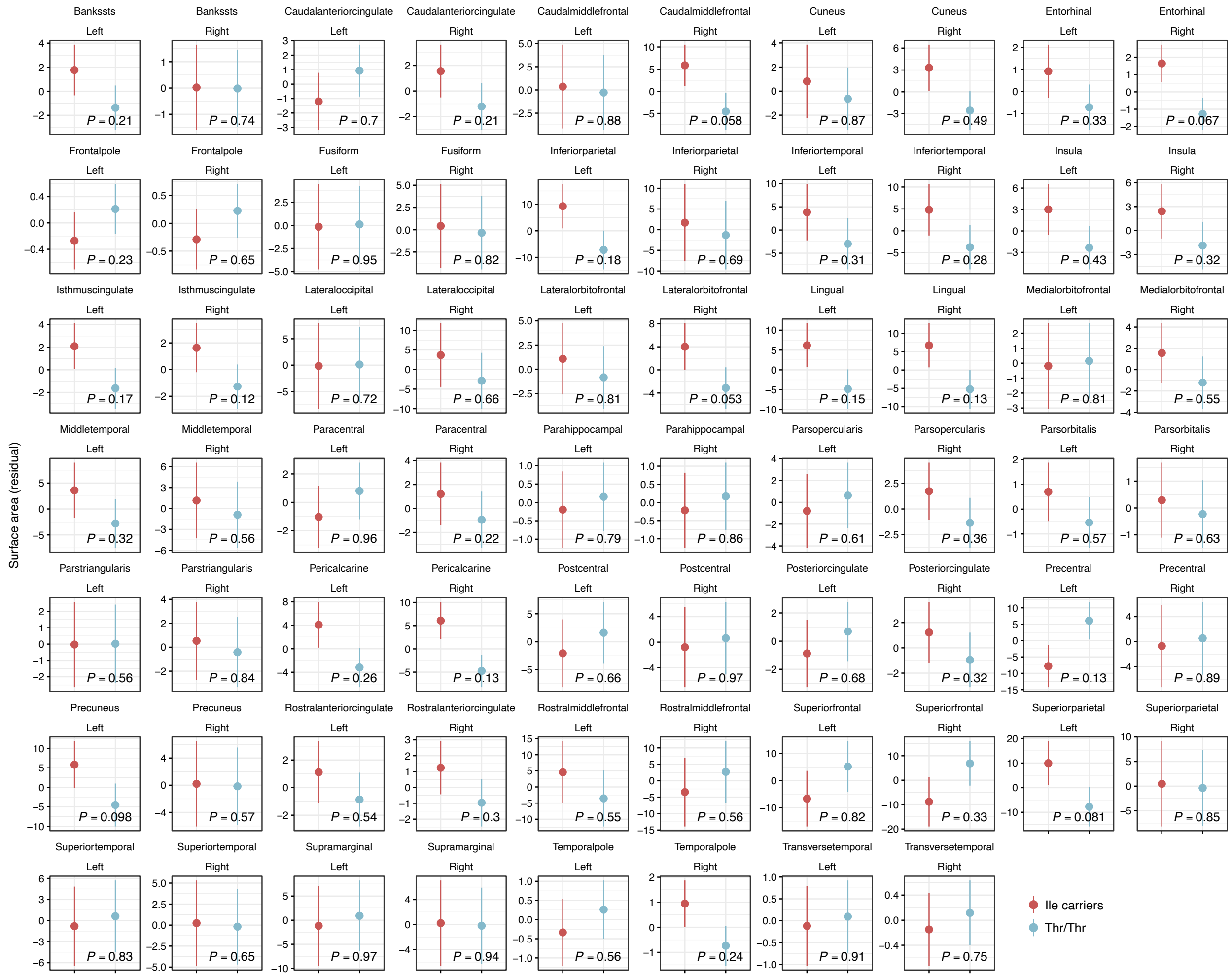


**Figure S1. Whole-brain comparison of regional cortical surface area between *VMAT1* genotype.** Residual cortical surface area values (adjusted for age, sex, and intracranial volume) are shown for each region of the Desikan–Killiany atlas in the left and right hemispheres. For each region, surface area was compared between *VMAT1* Thr/Thr homozygotes and Ile-carrier individuals using Wilcoxon's rank sum test. Points represent group means with error bars indicating standard errors. *P*-values from the Wilcoxon tests are displayed within each panel.

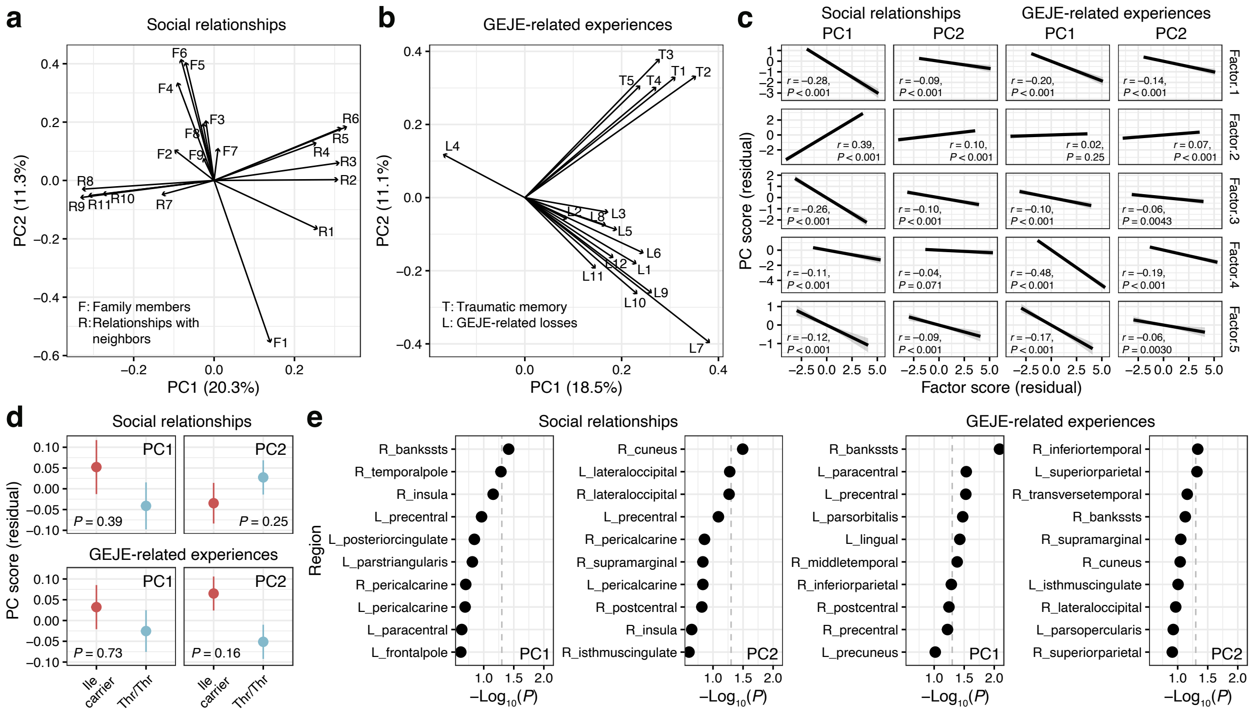


**Figure S2. Principal components of social relationships and GEJE-related experiences and their associations with psychological factors, *VMAT1* genotype, and cortical surface area.** **(a)** Biplot of the principal component analysis (PCA) conducted on social-relationship items, including family-relationship variables (F1–9) and neighborhood/social cohesion variables (R1–11). **(b)** Biplot of the PCA conducted on GEJE-related items, including variables regarding disaster-related exposures and losses (L1–12) and traumatic memory (T1–5). **(c)** Pearson correlations between lifestyle–experience PC scores and psychological factor scores (Factors 1–5), residualized for age and sex. **(d)** Distributions of PC scores by *VMAT1* genotype (Ile carriers vs. Thr/Thr). Points represent group means with error bars indicating standard errors. **(e)** Genotype-by-surface-area interaction tests predicting each PC score across all cortical regions. For each PC, linear models were fitted with the PC score as the dependent variable and *VMAT1* genotype, regional cortical surface area, and their interaction as predictors. Each point represents the *P*-value of the Genotype × SurfaceArea interaction term for one cortical region. Region labels are prefixed with L_ and R_ to denote the left and right hemispheres, respectively. The dashed gray line indicates $P = 0.05$.

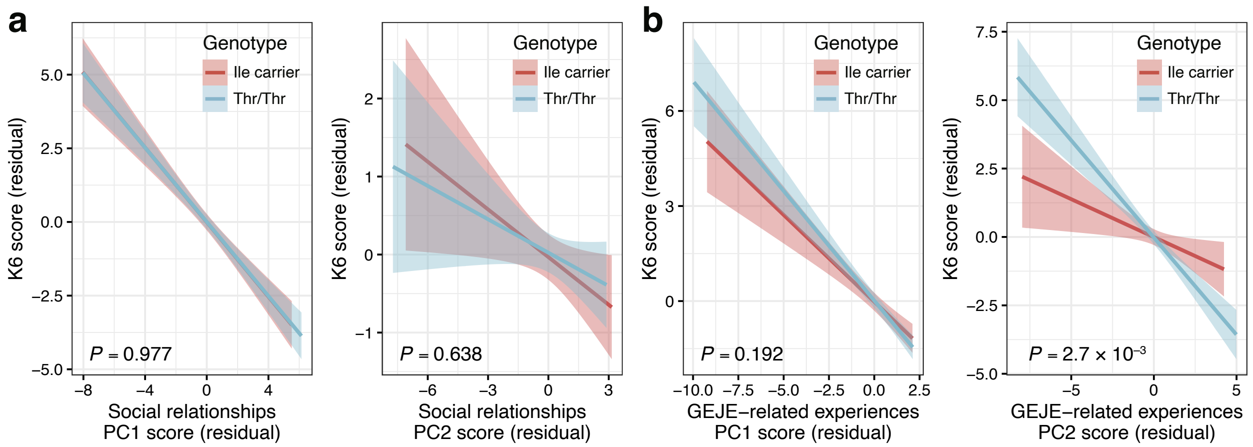


**Figure S3. *VMAT1* genotype-dependent effects of social and GEJE-related experiences on psychological distress (K6).** Relationship between age- and sex-corrected **(a)** social-relationship and **(b)** GEJE-related PC scores and K6 scores across genotypes. The *P*-value reflects the PC × genotype (Ile carrier vs. Thr/Thr) interaction term, indicating genotype-dependent effects of PC scores on K6. Shaded bands represent 95% CIs.

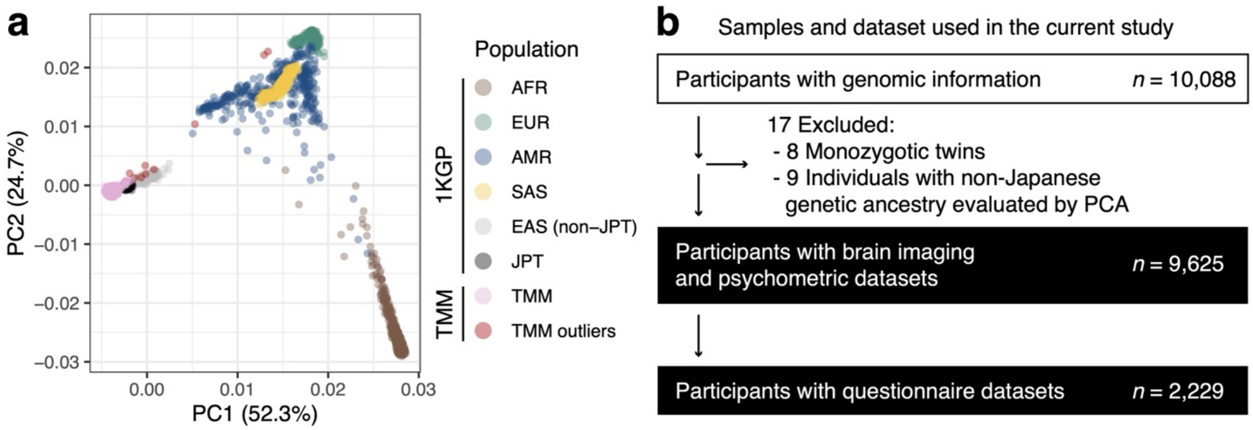


**Figure S4. PCA-based ancestry evaluation and sample selection flow. (a)** PCA of TMM participants (pink) merged with the 1000 Genomes Project reference panel. PC1 and PC2 capture continental-level population structure. Most TMM participants cluster within or adjacent to the JPT population, while outliers (dark red) were removed from downstream analyses. **(b)** Flow chart illustrating sample inclusion for genomic, MRI, psychometric, and questionnaire analyses. Among 10,088 genotyped participants, 17 individuals (8 monozygotic twins and 9 participants with non-Japanese genetic ancestry identified by PCA) were excluded, resulting in 9,625 individuals with MRI and psychometric data and 2,229 individuals with complete questionnaire data.
